# A lysine residue from an extracellular turret switches the ion preference in a Cav3 T-Type channel

**DOI:** 10.1101/2022.05.11.491449

**Authors:** W. Guan, K. G. Orellana, R.F. Stephens, B. Zhorov, J.D Spafford

## Abstract

A dominant sodium influx is mediated by a lysine residue in a novel extracellular loop of T-type Cav3 channels. Here we expressed calcium- and sodium-permeable splice variants which have exons 12b and 12a, respectively, as well as mutated channels containing exons Lys-12a-Ala and Ala-12b-Lys. We demonstrate that the mutant channels render high sodium permeability and calcium selectivity, respectively. Modelling illustrate that the pore lysine is salt-bridged to an aspartate residue immediately C-terminal to the second-domain glutamate of the selectivity-filter. We propose that a calcium ion chelated between the aspartate and the selectivity-filter glutamate is knocked-out by the incoming calcium ion in the process of calcium permeation, but sodium ions are repelled. The aspartate is neutralized by the lysine residue in the sodium-permeant variant, allowing for sodium permeation through the selectivity filter ring of four acidic residues akin to the prokaryotic sodium channels with four glutamates in the selectivity filter.

## Introduction

The distinct classes of voltage-gated calcium channels and sodium channels are defined by their selective passage of Ca^2+^ or Na^+^ (Hille, 2001). This ion selection is primarily determined by a pore selectivity filter contributed by an outer site composed of side chain residues consisting of DEKA (aspartate-glutamate-lysine-aspartate) in Na_V_ channels (Schlief et al., 1996) and a ring of five acidic residues, e.g. EE(D)EE or EE(D)DD, in the outer pore of Ca_V_ channels (Tang et al., 1993). The vital importance of the outer pore residues in defining ion selectivity has been demonstrated in the attribution of Ca^2+^ selectivity onto Na_V_ channels after glutamate residues were engineered into the Na_V_ channel selectivity filter (Heinemann et al., 1992). Many animal groups such as nematodes, parasitic platyhelminth (e.g. schistosomes), hemichordates and echinoderms completely lack voltage-gated Na_V_2 or Na_V_1 channel genes in their genomes (Fux et al., 2018). Even though nematodes don’t possess a Na_V_ channel gene, it is well established that nematodes generate cardiac-like action potentials in pharyngeal muscles that require a voltage-dependent Na^+^ current, but the gene responsible for this Na^+^ current has eluded discovery (Vinogradova et al., 2006, Franks et al., 2002). Similarly the giant pond snail, *Lymnaea stagnalis*, possesses cardiac action potentials requiring Na^+^ and Ca^2+^, but its singleton Na_V_1 sodium channel gene in its nervous system doesn’t express in the snail cardiovascular system (Senatore et al., 2014). We have demonstrated that the source of the Na^+^ current in the snail heart is a mibefradil and nickel sensitive low voltage-gated Cav3 T-type current (Senatore et al., 2014). We can generate the high Na^+^ passing T-type current from the snail heart in the expression of a unique, alternatively-spliced isoform of the LCav3 T-type channel gene containing exon 12a expressed in the snail heart (Senatore et al., 2014). Exon 12b, which doesn’t express in the snail heart, but predominates in skeletal muscle tissue, confers the typical phenotype of mostly Ca^2+^ passing currents onto the LCav3 T-type channel (Senatore et al., 2014). We demonstrate that the human calcium-selective Cav3.2 channel can be converted into T-type calcium channel with a preference for passage of Na^+^ over Ca^2+^, in chimeras which includes the swapping of exon 12a from the snail LCa_V_3 T-type channel (Guan et al., 2020). Exons 12a and 12b code for the L5 extracellular turret in Domain II (IIS5-P1) of the four domain Cav3 T-type channel, terminating just upstream of the hallmark pore selectivity filter residues known for conferring the characteristic ion selectivity of Ca_V_ and Na_V_ channels (Senatore et al., 2014, Guan et al., 2020). What has eluded discovery is an atomic model to explain how a typically fast, Ca^2+^-selective T-type channel converts into a kinetically slow ion channel with a preference for passage of Na^+^ over Ca^2+^ without alterations to the hallmark pore selectivity filter or amino acids in the voltage sensor domain. Here, we illustrate that what varies is the presence of a single, lysine residue within exon 12a, but not exon 12b to neutralize a critical aspartate residue in the outer pore of Domain II contributing to calcium selectivity. Appearance of a short, alternative extracellular turret evolved to infiltrate and adjust the pore’s ion permeability, provides an ingenious and economical way for nematodes and mollusks to generate multiple phenotypes resembling multiple Ca_V_ and Na_V_ channels within their singleton T-type channel in the absence of expression of differing Ca_V_ or Na_V_ channel isoforms.

## Results and Discussion

A first experimentally derived structure of a Ca_V_3 T-type channel by cryo-electron microscopy at < 3.4 A° resolution, reveals a conservation with other Ca_V_1 and Ca_V_2 channels in the largest extracellular loops IS5-P1 and IIIS5-P1, rising and looming over the pore selectivity filter below, stabilized by many conserved disulfide bonds (Zhao et al., 2019). Domain IV contains the longest extracellular loop IVP2-S6 (Zhao et al., 2019), where protostome invertebrates Cav3 T-type channels vary from vertebrate Cav3 T-type channels in possessing a longer loop containing an extra intra-turret cysteine bridge (Guan et al., 2020). The shortest is the IIS5-P1 extracellular loop, where Ca_V_3 T-type channels strikingly vary between vertebrates and invertebrates, and where Cav3 T-type channels differ more dramatically from other Ca_V_1 and Ca_V_2 channels (Senatore et al., 2014). In the cryo-EM structure of human Cav3.1 channel a cysteine in the IIS5-P1 loop forms a disulfide bridge to a cysteine in the extracellular loop (IS1-S2) of the voltage sensor domain (**Fig. 1**) (Zhao et al., 2019). This unique tethering of the voltage-sensor and pore domains is likely to contribute to the reported redox sensitivity and variability in ion channel kinetics of Ca_V_3 T-type channels (Todorovic et al., 2001, Karmazinova et al., 2010).

**Figure 1.**
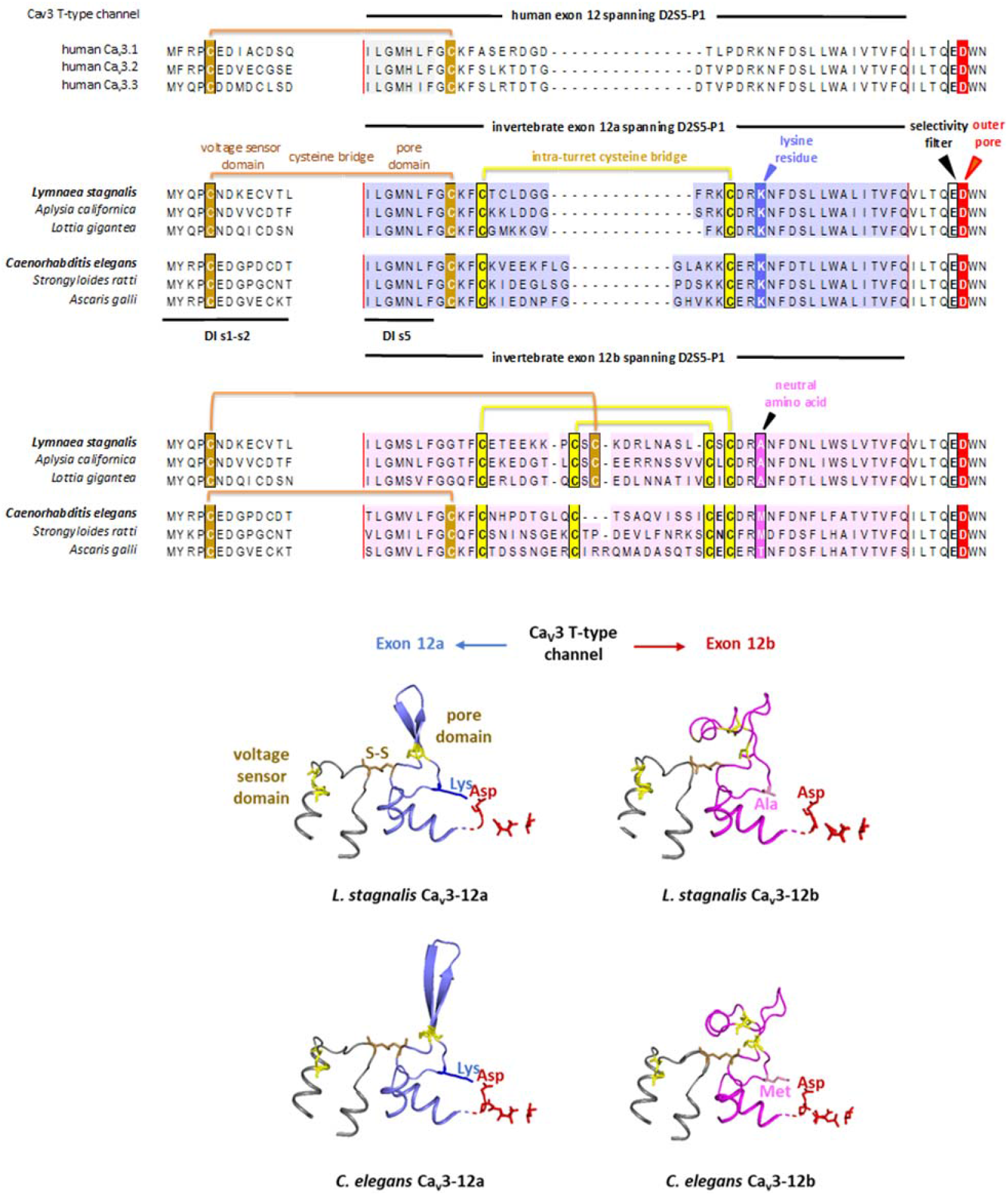
Alternative exon 12a and exon 12b (delimited by exon borders illustrated by red lines, top panel) are indicated for molluscan and nematode Cav3 T-type channels, where the positively-charged lysine residue in exon A (blue color) but not the neutral amino acid (A, M, T) in exon B lies adjacent to the outer pore aspartate residue (bottom panel) to neutralize and create sodium permeant Cav3 T-type channels. Locations of the cysteine bridge spanning the voltage sensor domain and the pore domain are indicated for the molluscan exon 12a isoform and nematode exon 12b isoform. (Top panel) Aligned amino acid sequences for human (Cav3.1, Cav3.2 and Cav3.3) and singleton homologs of Cav3 T-type channels from representative mollusks (*Lymnaea stagnalis, Aplysia californica* and *Lottia gigantea*) and nematodes (*Caenorhabditis elegans, Strongyloides ratti* and *Ascaris galli*). (Bottom panel) AlphaFold2 generated models illustrating mollusk, *L. stagnalis* and nematode, *C. elegans*. Illustrated are the regions spanning the extracellular loop spanning membrane segments s1-s2 in Domain I of the voltage sensor and the pore domain spanning exon 12 from the end of membrane segment s5 to the extracellular DIIL5 turret terminating upstream of the re-entrant pore helix 1 and the selectivity filter residue (glutamate, E) followed by the outer pore (aspartate, D) located above the selectivity filter.

Protostome invertebrates from flatworms, arthropods, mollusks to annelids contain alternative IIS5-P1 extracellular loops encoded by exon 12a and exon 12b (Guan et al., 2020, Senatore et al., 2014). AlphaFold2 generated structures indicate that exon 12a within protostome Ca_V_3 channels possess a similarly placed cysteine forming a disulfide bond to the voltage sensor domain like in the mammalian Ca_V_3.1 channel (Zhao et al., 2019) (**Fig. 1**). The cysteine linking to the voltage-sensor domain of exon 12b is contained within a pair of intra-turret cysteine bridges contained in exon 12b of most protostome invertebrates such as in the mollusk, *Lymnaea stagnalis* rather than outside the intra-turret cysteine bridge of exon 12a (**Fig. 1**). The exception is for nematode Cav3 channels, such as *Caenorhabditis elegans* where the cysteine bonded to the voltage sensor domain remains in the same position for exon 12a and exon 12b outside the one or two pairs of intra-turret cysteine bridges, respectively (Guan et al., 2020, Senatore et al., 2014) (**Fig. 1**). Notably we find that all measured kinetic features (activation, inactivation, deactivation, recovery from inactivation) to a significant degree, are slowed in molluscan LCa_V_3 channels containing exon 12a compared to the faster kinetics with exon 12b which possesses the cysteine bridging to the voltage sensor domain localized within the double set of intra-turret cysteines (**Table I**).

**Table 1:**
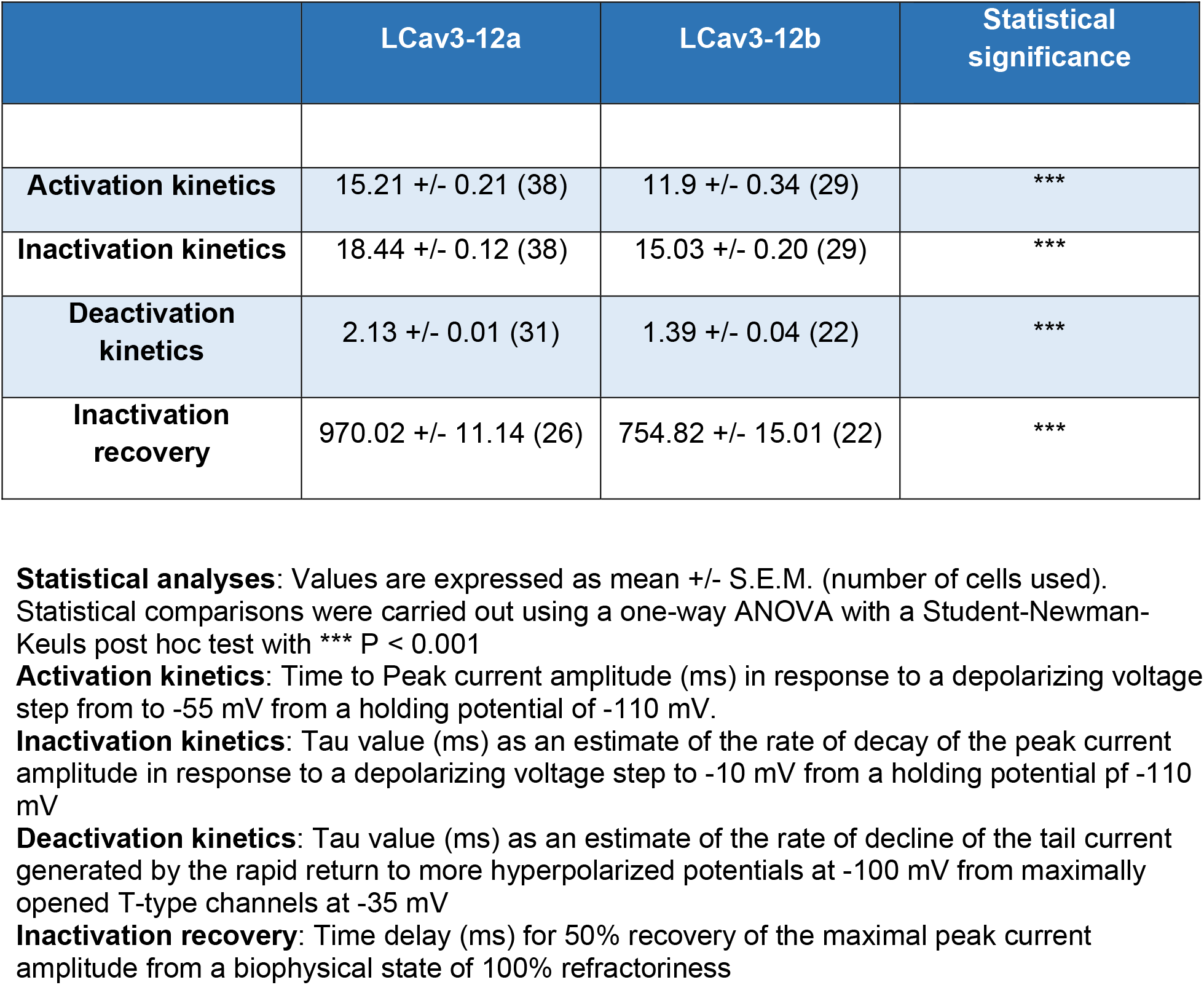
Comparison of the biophysical parameters of snail LCa_v_3 channels expressed with exon 12a or exon 12b in HEK-293T cells.

Exons 12a and exons 12b also specifically vary in one other residue within all molluscan (e.g. *Lymnaea stagnalis*) (Senatore et al., 2014) and nematode (e.g. *Caenorhabditis elegans*) (Steger et al., 2005) Cav3 channels. This is a positively-charged lysine residue within exon 12a (K1067, Genbank Acc. # JX292155 and K819, Genbank Acc. # AAP84337, respectively) compared to a neutral alanine or methionine residue at a similar position within exon 12b (A1078, Genbank Acc. # AAO83843 and M826, Genbank Acc. # AAP79881, respectively). Despite differing sequences and number of intra-turret cysteine bridges between exon 12a and exon 12b, AlphaFold2 derived structures overlay into an identical positioning of the neutral amino acid in exon 12b and the charged lysine in exon 12a, where the protonated side chain lysyl residue extends to approach within 2.6 angstroms of the negatively charged aspartate residue found in similar position of the outer pore within all calcium (Ca_V_1, Ca_V_2, and Ca_v_3) channels.

To assess the relative sodium and calcium permeation through molluscan LCa_V_3 channels, we measured the relative fold increase in peak current amplitude when external Na^+^ replaces weakly permeant cation, NMDG^+^ in the presence of external Ca^2+^. Direct mutation of the outer pore’s aspartate residue to a more neutral asparagine residue (D1087N) imitates the consequence of neutralizing the outer pore’s aspartate residue by juxtaposition of the lysine residue from the IIS5-P1 extracellular turret (**Fig. 2a**), leading to a rise in the relative Na^+^ permeation through LCav3-12b D1087N mutated channels compared to wild-type LCav3-12b channels (**Fig. 2b, 2c**). We confirm the identical placement of the neutral alanine residue of exon 12b and the positively-charged lysine of exon 12a in the IIS5-P1 extracellular turret in the computationally-derived atomic models, by analyzing the consequences of swapping the positively-charged lysine residue of exon 12a for the neutral alanine residue in exon 12b. As predicted, the swapped amino acid of LCa_V_3-12a K1067A generates the Ca^2+^ selective phenotype resembling LCa_V_3-12b within LCa_V_3-12a. And, vice-versa, LCa_V_3-12b A1078K mutated channels resembles the phenotype of the highly Na^+^ permeant, LCa_V_3-12a channels. While the selective mutational change to a lysine or alanine residue in exon 12 conferred a dramatic change in the preference for the passage of Na^+^ and Ca^2+^, respectively, the mutated LCa_V_3-12a and LCa_V_3-12b channels were still distinguishable by the characteristically slower kinetics of LCa_V_3-12a channels compared to the faster kinetics of LCa_V_3-12b channels (exemplified by deactivation kinetics in **Fig. 2c**).

**Figure 2.**
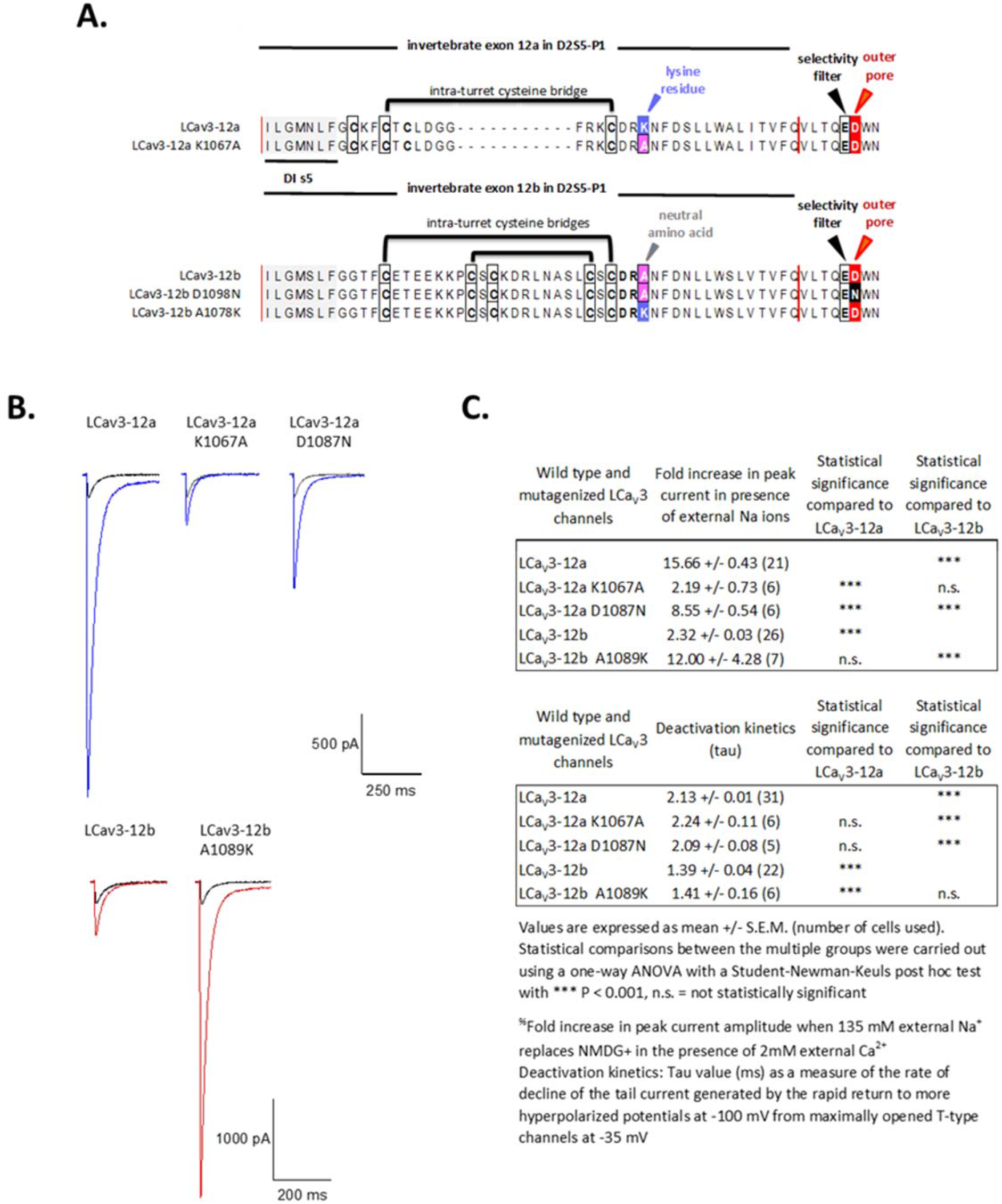
Mutational analyses illustrating the intra-convertibility of exon 12a and exon 12b within molluscan LCav3 channels in generating highly sodium permeant Cav3 T-type channels with a lysine residue in homologous position within the D2L5 extracellular turret. (A) Aligned amino acid sequences indicating the single amino acid swap (lysine vs alanine) in D2L5 extracellular turret with mutants, LCav3-12a K1067A and LCav3-12b A1089K. The consequence of the lysine residue in neutralizing the calcium beacon residue is mimicked in the generation of the LCav3-12a D1987N mutant. (B) Representative ionic current traces of LCav3 channels transfected in HEK-293T cells and recorded using whole cell voltage clamp. Black current traces were recorded in the presence of external Ca2+ and weakly permeant cation (NMDG+), and the red and blue colored traces were recorded after replacement of NMDG+ equimolar Na+. (C) Table of statistics of wild type and mutagenized LCav3 channels.

### A short IIS5-P1 loop evolved to co-opt a Cav3 T-type channels into a slow Na_V_ channel phenotype without altering the pore’s selectivity filter

We demonstrate here a first example of how nature has engineered a means for an ion channel to switch its phenotype with the positioning of a single glutamate residue in an alternative exon to generate a switch in preference for passage of Na^+^ over Ca^2+^, utilizing a short alternative extracellular L5 turret in Domain II spanning less than 1% of the massive ~ 322 kDa LCa_V_3 T-type channel.

Despite the divergence in their sequences and differing one and two intra-turret cysteine bridges in exon 12a and exon 12b, respectively, the IIS5-P1 extracellular turret from both exons structurally overlap to arrive at a positively charged lysine contained in exon 12a, in identical position of the neutral alanine of exon 12b (**Fig. 3a**). Notably, the lysine in exon 12a is salt-bridged to the second-domain aspartate in the filter EE(D)DD, but not the neutral alanine of exon 12b (**Fig. 3a**).

**Figure 3.**
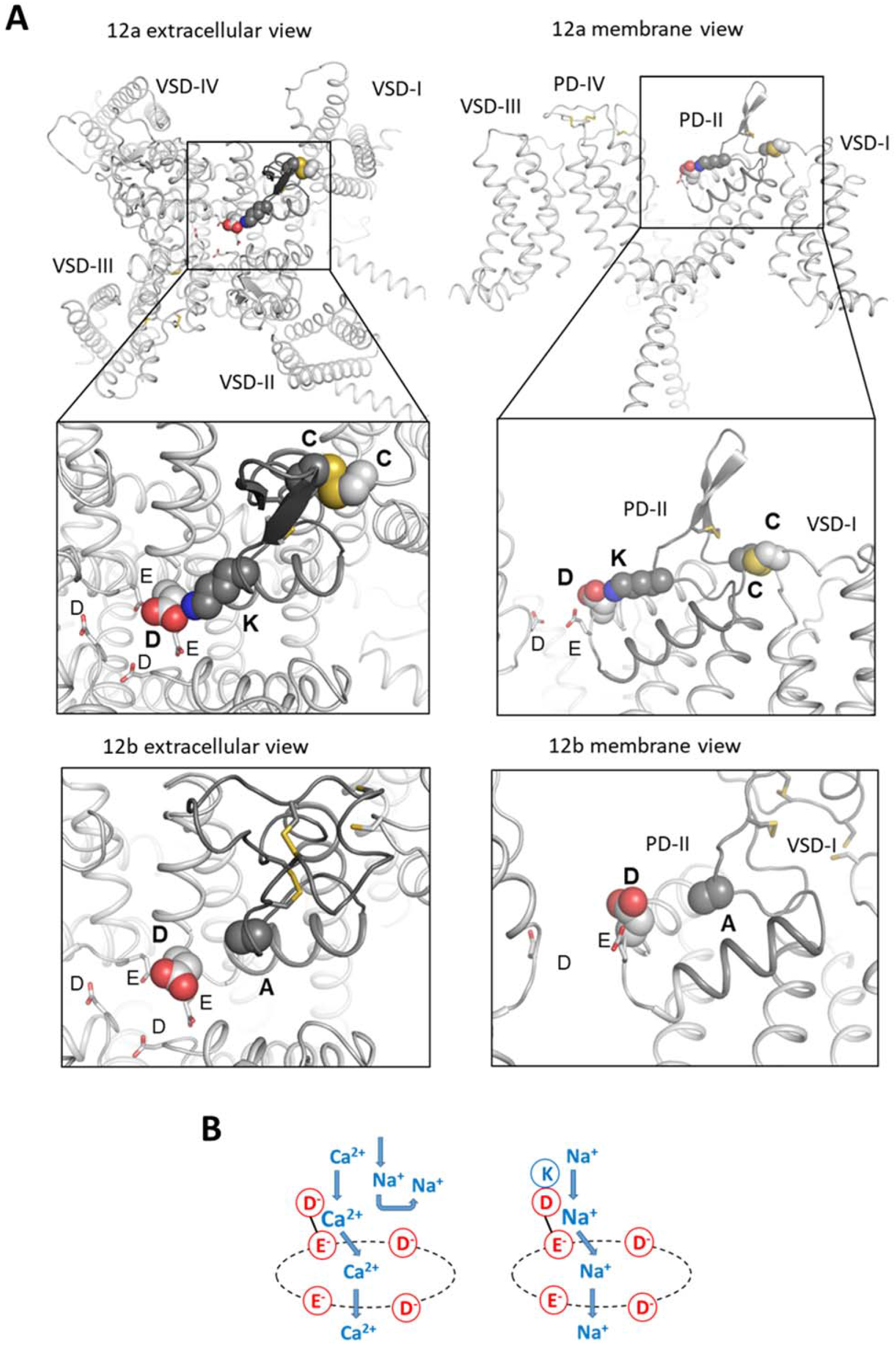
**(A) AlphaFold2 models of LCav3 T-type channel.** Exon 12a (dark) has a lysine salt-bridged to the aspartate, which is immediately C-terminal to the Domain II glutamate in the EEDD selectivity-filter ring. An alanine in exon 12b (dark) is in the position of the lysine in exon 12a. **(B) Possible mechanism for the different ion selectivity phenotypes of LCav3 channels with exons 12a and 12b.** Ca^2+^ ions are chelated between carboxylate groups in the dipeptide ED of Domain II in LCav3-12b. An incoming Ca^2+^ ions knocks-out the chelated Ca^2+^, which moves down into the inner pore. An incoming Na+ ion is repelled by the chelated Ca^2+^. In the channel with exon 12a, the lysine residue from the exon is salt-bridged to the ED and neutralizes its negative charge. The ring of four acidic residues would be sodium-permeable like the prokaryotic sodium channels with EEEE residues in selectivity-filter rings.

We confirm the atomic modelling that swapping the neutral alanine with the positively-charged lysine of the extracellular exon 12a turret onto exon 12b, generates a LCa_V_3 channel with a preference for passage of Na^+^ over Ca^2+^, reminiscent of LCav3-12a, while the reverse (lysine to alanine) mutation in the extracellular D2L5 turret generates a more traditional Ca^2+^-selective Cav3 channel onto LCav3-12a, reminiscent of LCav3-12b.

The atomic models suggest the following mechanisms for the switch in preference for passing Ca^2+^ or Na^+^ in the Cav3 channels with exons 12a and 12b (**Fig. 3b**). In LCav3 with exon 12a, Ca^2+^ is chelated between carboxylate groups in the dipeptide ED of the Domain II pore. An incoming Ca^2+^ knocks-out the chelated Ca^2+^, which moves down into the inner pore while an incoming Na^+^ is repelled by the chelated Ca^2+^. In LCav3 with exon 12b, the lysine residue from the exon is salt-bridged to the ED and neutralizes its negative charge. The selectivity-filter ring with the net negative charge of −4 would be sodium-permeable like prokaryotic sodium channels with the EEEE residues in the selectivity-filter ring (Chakrabarti et al., 2013).

We illustrate that partial neutralization of the aspartate (D) by site specific mutation to an asparagine (N) in the outer pore, increases the preference for passage of Na^+^ over Ca^2+^ in a manner consistent by the atomic modelling prediction of the charge nullification of the aspartate by the juxtaposition of the positively charged lysine residue next to aspartate residue in the extracellular D2L5 turret within exon 12a of the molluscan Cav3 T-type channel.

### Support from physiological experiments which demonstrate a change in ion selectivity from Ca to Na channels in the neutralization of the outer pore aspartate with the turret lysine

The charge nullification of the outer pore’s aspartate by the juxta positioning of a critically-positioned lysine instead of an alanine in homologous position of the extracellular turret within exon 12, is consistent with our observations of a changing ion selectivity phenotype from Ca^2+^ to Na^+^ in exon 12a containing LCa_V_3 channels. We have analyzed the relative changes in Ca^2+^ and Na^+^ flux due to exon 12b and exon 12a in chimeras to engineer a Na^+^-selective T-type channel within a human Ca_V_3.2 T-type channel background, and to engineer a Ca^2+^-selective T-type channel with the human extracellular D2L5 turret onto a snail LCa_V_3 channel background (Guan et al., 2020). This is reflected in an increased contribution of the inward Na^+^ current estimated as the fold change in peak inward cation current size in physiological external Ca^2+^ concentration, when an equimolar quantity of external Na^+^ replaces weakly permeant monovalent ion, NMDG^+^ (N-methyl-D-glucamine). The changing relative permeability of the Na^+^ flux compared to Ca^2+^ flux due to exon 12a can be estimated in bi-ionic reversal experiments as the degree of reversal potential change in response to Ca^2+^ influx to a competing efflux of different monovalent ions (Li^+^, Na^+^, K^+^, Cs^+^), in response to high concentrations of monovalent ions in internal solutions instead of external solutions (Senatore et al., 2014, Guan et al., 2020). Traditional Ca^2+^-selective channels generate a “*U-shaped*” response curve in response to increasing doses of calcium ions in the presence of external Na^+^ (Senatore et al., 2014, Guan et al., 2020). The U-shaped response to increasing Ca^2+^ is considered a reflection that Na^+^ ions are unable to compete for the limited cation binding sites as Ca^2+^ effectively repels the funneling of Na^+^ through the Ca^2+^-selective pore in the presence of low 10 μM external Ca^2+^ concentrations, and where Ca^2+^ influx dominates Na^+^ influx at physiological (mM) levels of external Ca^2+^ through calcium-selective channels (Senatore et al., 2014, Guan et al., 2020).

The property of a high Na^+^ selectivity over Ca^2+^ in the molluscan LCa_V_3 channel pore containing exon 12a is reflected in the transformation of the “*U-shaped*” response curve in response to increasing external Ca^2+^ into a “*reverse S*” response curve, where low 10 μM external Ca^2+^ concentration is ineffective in the blocking of the Na^+^ influx, and rises in external Ca^2+^ to external physiological (mM) concentrations does not lead to greater Ca^2+^ passage through the Na^+^ selective T-type channel pore, but rather there is an observed increasing block of the T-type channel current because Ca^2+^ is not capable of permeating through the sodium-selective, LCav3-12a T-type channel pore at physiological (mM) concentrations of Ca^2+^ (Senatore et al., 2014, Guan et al., 2020).

### All known mollusks and nematode Cav3 T-type channel transcripts possess predicted T-type sodium channels expressed with exon 12a

All available nematodes sequences surveyed in GenBank, include model organism, *C. elegans*, possess AlphaFold2 generated atomic structures with the charge nullifying, lysine residue adjacent to the outer pore aspartate, to generate a mostly Na^+^-passing T-type channel with their exon 12a like in mollusks (Senatore et al., 2014, Guan et al., 2020). All exon 12b sequences of nematodes possess a neutral residue (alanine, methionine, serine, and threonine) in identical position to the lysine residue to generate a mostly Ca^2+^ passing T-type channel (Senatore et al., 2014, Guan et al., 2020). The only notable structural difference between molluscan and nematode T-type channel with exon 12a and exon 12b is a different location of the cysteine that bridges from D2L5-P1 extracellular turret to the cysteine in the voltage sensor domain. The cysteine bridging the voltage sensor is within rather than outside the two pairs of intra-turret cysteines in all prostostome invertebrates outside of nematodes (**Fig. 1**).

### The differing turret cysteine bridging to the voltage-sensor domain in exon 12a generates a kinetically slower T-type sodium channel appropriate for the molluscan heart

The differing location of the cysteine tethered to the voltage sensor domain within extracellular loops of exon 12a and exon 12b, might be a reflection of the differing cellular conditions requiring kinetically fast or slow Cav3 T-type channels within protostome invertebrates. In the case of mollusks, they possess a traditionally fast, Na^+^-selective Na_V_1 channel, but this gene is limited in expression to the nervous system (Senatore et al., 2014). A Na^+^-permeant T-type channel with exon 12a is the only isoform of T-type channel expressed in the molluscan heart, where it serves as a surrogate Na^+^ current in the absence of expression of the singleton Na_V_1 channel in mollusks (Senatore et al., 2014). Exon 12a imparts a dramatically slowing of its activation, inactivation, deactivation, and recovery from inactivation in LCa_V_3 T-type channels compared to exon 12b. And this slowing of kinetics suits the requirements of a surrogate Na^+^ current in the snail heart, akin to a heart specific isoform in vertebrates, such as Na_V_1.5 (Wang et al., 1996), to establish a cardiac action potential with a prolonged repolarization phase required for refilling of the heart, with a built in refractoriness to prevent ectopic heart beats.

While the co-opting of a kinetically slower, Na^+^-permeant T-type channel serves the purposes of mollusks *in lieu* of expression of their Na_V_1 channel gene within molluscan hearts, *C. elegans* lack expression of any Na_V_2 or Na_V_1 sodium channel gene (Bargmann, 1998). Nematode Ca_V_3 T-type channel sequences generate precisely aligned extracellular turret structures in AlphaFold2 with identical positioning of the outer pore aspartate nullifying lysine residue with exon 12a and a precisely identical positioning for a neutral residue in exon 12b. But there is also a difference in nematode Ca_V_3 channels where the cysteine linkage to the voltage sensor from the D2S5-P1 extracellular loop is outside the one and two intra-turret cysteine bridges for both exons 12a and exon 12b. The unique configuration of exon 12a and exon 12b within nematodes provide an optional sodium permeability for the nematode Ca_V_3 channel to meet expectations of action potential spike generation such as in the pharynx of *C. elegans*, where voltage-gated sodium currents have been observed (Vinogradova et al., 2006, Franks et al., 2002).

## Conclusions

The Ca_V_3 channels differ from other (Ca_V_1 and Ca_V_2) calcium channels and sodium channels in the diversity of cysteine bridges within the shortest of the four S5-P1 extracellular turrets in Domain II, and in the largest of the P2-S6 extracellular turrets contained in Domain IV (Stephens et al., 2015, Zhao et al., 2019). The competing cysteine tethers resemble a “*tug of war*” at extracellular turrets. One cysteine tether link the pore domain to the voltage sensor domain by a disulphide bond to impart kinetic changes in molluscan Cav3 T-type channels. One and two unique intra-turret cysteine bridges are also present within exon 12a and exon12b spanning D2S5-P1 extracellular turrets of protostome invertebrates, respectively. The intra-turret disulphides are likely associated with providing the structural integrity of the pore necessary to impart the Na^+^ and Ca^2+^ selectivity changes onto Cav3 T-type channels (Guan et al., 2020). Cysteine to alanine mutations which disrupt the intra-turret disulfide bridges on exon 12a and exon 12b not only dramatically influences the relative Ca^2+^ and Na^+^ selectivity through LCa_V_3 channels but also dramatically alter the potency of Zn^2+^ and Ni^2+^ block by ~ 50× and ~10× fold respectively, including the loss of the slowing of inactivation kinetics during Zn2+ block (Guan et al., 2020).

We also observe an additional, intra-turret cysteine bridge within P2-S6 extracellular loop of Domain IV within all protostome invertebrates, balancing the reinforcement of extra cysteine bridges within their S5-P1 extracellular turrets in across from the Ca_V_3 T-type channel pore in Domain II. We have illustrated that the extra cysteine bridge in D4P2-S6 extracellular turret of Domain IV is necessary for the D2S5-P1 extracellular turret contained in exon 12a from mollusks to impart the characteristically high Na^+^ preference over Ca^2+^ onto human Ca_V_3.2 channel pores (Guan et al., 2020). Increases in cysteine bridge number over the basic complement of other Ca_V_1 and Ca_V_2 channels within extracellular turrets, first appears in the basal group of extant animals containing “*true*” nervous systems, the cnidarians, where there are two different Ca_V_3 T-type channel genes in anthozoans and scyphozoan cnidarians, one resembling the vertebrate condition with a Cav3 T-type gene coding for one cysteine bridge in D4P2-S6 extracellular loop and another Cav3 T-type gene resembling the protostome invertebrate condition with a gene coding for two cysteine bridges in D4P2-S6 extracellular loop (Guan et al., 2020). The hotbed of differing configurations of cysteine bridges within extracellular turrets provides a capacity to differentially modulate the functional diversity of T-type channels indirectly, where the extracellular turrets are tethered to induce functional changes indirectly, akin to “*marionette strings*” with cysteine-enriched rich extracellular loops imparting changes to ion channel kinetics and ion selectivity without specific amino acid mutation of the voltage sensor domain or the T-type channel pore’s ion selectivity filter, respectively, *per se*.

## Materlals and Methods

### Reagents and Tools Table

| REAGENT or RESOURCE | SOURCE | IDENTIFIER |
| --- | --- | --- |
| **Chemicals, Peptides and Recombinant Proteins** |  |  |
| Dulbecco's modified Eagle's medium | Millipore Sigma-Aldrich | Cat # D5796 |
| sodium pyruvate | Millipore Sigma-Aldrich | Cat # S8636 |
| fetal bovine serum | Wisent Bioproducts | Cat # 080-910 |
| Penicillin (10,000 U/mL)<br>Streptomycin (10 mg/mL) | Millipore Sigma-Aldrich | Cat # P4333 |
| Trypsin-EDTA solution | Millipore Sigma-Aldrich | Cat # T4049 |
| Poly-L-lysine solution | Millipore Sigma-Aldrich | Cat # P4707 |
| HEPES | Millipore Sigma-Aldrich | Cat # 391338 |
| **Experimental Models: Cell Lines** |  |  |
| HEK-293T | ATCC | Cat # CRL-3216 |
| **Recombinant DNA** |  |  |
| LCav3-12a (Cav3 T-type channel from *Lymnaea stagnalis* with configuration + exon 8b, + exon 12a, - exon 25C) | Senatore *et al.* 2010 | RRID: Addgene_ |
| LCav3-12b (Cav3 T-type channel from *Lymnaea stagnalis* with configuration + exon 8b, + exon 12b, - exon 25C) | Senatore *et al.* 2014 | RRID: Addgene_ |
| LCav3-12a K1067A | this manuscript | RRID: Addgene_ |
| LCav3-12a D1087N | this manuscript | RRID: Addgene_ |
| LCav3-12b A1089K | this manuscript | RRID: Addgene_ |
| **Software and Algorithms** |  |  |
| pClamp10 | Molecular Devices | https://www.moleculardevices.com/products/axon-patch-clamp-system/acquisition-and-analysis-software/pclamp-software-suite |
| Origin-Pro 2017 | Originlab | https://www.originlab.com/origin |
| AlphaFold v2.0 | Google | https://github.com/deepmind/alphafold |
| PYMOL 2.5 | Schrödinger | https://pymol.org/2/ |
| ChimeraX | Resource for Biocomputing, Visualization, and Informatics at UCSF | https://www.rbvi.ucsf.edu/chimerax/ |

## METHODS AND PROTOCOLS

Further information and requests for resources and reagents should be directed to and will be fulfilled by the Lead Contact, J. David Spafford.

### Materials availability statement

Addgene catalog numbers are given in Key Resources table and additional constructs will be deposited in Addgene.

## EXPERIMENTAL MODEL DETAILS

### HEK-293T cell line

Human embryonic kidney (HEK) 293T cell lines (M. Calos, Stanford University) were plated onto cell culture flasks and coverslips coated with poly-L-lysine, and cultured in a 5% CO2 incubator at 37°C. HEK-293T cells were cultured in Dulbecco’s modified Eagle’s medium (Millipore Sigma-Aldrich) with 10% fetal bovine serum (Millipore Sigma-Aldrich) and supplemented with 0.5% (v/v) penicillin-streptomycin solution (Millipore Sigma-Aldrich).

## METHOD DETAILS

### Molecular biology

The mostly Ca2+ passing isoform of the invertebrate LCa_V_3 T-type channel (GenBank Accession #: AAO83843), isolated from pond snail, *Lymnaea stagnalis*, was expressed and characterized in a configuration contained exon 12b, as well as optional exon 8b spanning the I–II linker, but lacking exon 25c of the III–IV linker (Senatore and Spafford, 2010). Novel exon 12a isoform (+ 8b, − 25C) deposited as GenBank Accession # AFN89594, is compared with exon 12b isoform (+ 8b, − 25C) which is the configuration of the three exons that is more commonly expressed in the snail brain than in the snail heart, where there is exclusive expression of the mostly Na^+^ current passing Ca_V_3 T-type channel with exon 12a (Senatore et al., 2014).

Site-directed mutations within exon 12 of full length Cav3 channels (LCav3-12a K1067A, LCav3-12a D1087N, LCav3-12b A1089K) were generated by swapping synthetic gene fragments containing the site-specific mutations (ordered from BioBasic Canada) spanning novel silent restriction sites AvrII and Eco47III (AfeI) introduced at positions: 3,338 to 3,567 spanning the coding sequence for exon 12a (39 amino acids) and exon 12b (50 amino acids). Synthetic DNA (ordered from BioBasic Canada) spanning the AvrII and Eco47III restriction sites were inserted into a subclone in pGEMT vector at unique BglII and SalI restriction sites (positions: 1,391 to 4,521). Successfully ligated subclones containing the site directed mutations, were reintroduced into the full length LCav3 cDNA cloned in pIRES2-EGFP at the BglII and SalI restriction sites. LCav3-12a K1067A, LCav3-12a D1087N, LCav3-12b A1089K contained a unique BspEI, PvuI and KpnI silent restriction sites, respectively, for rapid validation of individual cloned plasmid stocks for their unique L5_II_ extracellular loop identity in LCa_V_3.

### Molecular modelling

Molecular models of amino acid sequences were generated using AlphaFold2 derived from molluscan Cav3 T-type channel from *Lymnaea stagnalis* containing exon 12b (Genbank Acc. # AAO83843) or containing exon 12a (Genbank Acc. # JX292155), and from nematode Cav3 T-type channel from *Caenorhabditis elegans* containing exon 12b (Genbank Acc. # AAP79881.1) or containing exon 12a (Genbank Acc. # AAP84337.1). Structural models were overlaid and curated in ChimeraX (Resource for Biocomputing, Visualization, and Informatics at UCSF) or PyMOL 2.5 (Schrödinger)

### Cell line transfection

LCav3 clones including mutated derivatives were expressed within bi-cistronic, mammalian expression, pIRES2-EGFP vector, which generates green fluorescence upon mercury lamp excitation for identifying positively transfected HEK-293T cells. Six μg of LCav3 channel containing plasmids were heterologously expressed, by calcium-phosphate transfection into human embryonic kidney cell line (HEK293T, M. Calos, Stanford University) at 40–50% confluency. After overnight transfection, the cells were washed three times with culture media and incubated at 28°C in a humidified, 5% CO2 chamber for three to five days. Cells were replated in 60 mm (diameter) sterile Petri dishes containing eight poly-lysine coated glass coverslips (Circles No. 1-0.13 to 0.17 mm thick; Size: 12 mm), (ThermoFisher Scientific) and allowed to recover at 37°C for four hours then left at 28°C for at least three days before patching. Complete details of our methods for optimized transfection and recording of ion channels is available as a video journal entry at: http://www.jove.com/index/Details.stp?ID=2314 (Senatore et al., 2011).

### Electrophysiology

Calcium currents through LCav3 T-type channels was recorded in the presence in external calcium containing extracellular recording solution consisting of (in mM): 2 CaCl_2_, 135 NMDG^+^ (N-Methyl-D-Glucamine); 25 TEA (Tetraethylammonium), 10 HEPES (4-(2-hydroxyethyl)-1-piperazineethanesulfonic acid), at a pH of 7.4 titred with TEA-OH. The evaluation of the relative sodium conductance through LCav3 T-type channels were evaluated in the presence of the external calcium-containing extracellular recording solution described above where 135 mM Na^+^ replaced 135 mM NMDG^+^.

Whole cell electrophysiology recordings were obtained using an Axopatch 200B or Multiclamp 700B amplifiers (Molecular Devices), sampled through a Digidata 1440a A/D converter (Molecular Devices) to a PC computer. Patch pipettes for recording transfected HEK-293T cells had pipette resistances of 2.5 to 5 MΩ, and with typical access resistance maintained after breakthrough between 2.5 MΩ and 6 MΩ (HEK-293 T cells). Internal recording solution consisted of (in mM): 110 mm CsCl, 10 EGTA, 3 Mg-ATP mm CaCl2, 0.3 mm Tris-GTP, and 10 HEPES, pH 7.2, at a pH of 7.2 with CsOH.

Series resistance was compensated to 70% (prediction and correction; 10-μs time lag). Offline leak subtraction was carried out using the ClampFit 10.2 software (Molecular Devices). For all recordings, offline leak subtraction was carried out and data was filtered using a 500 Hz Gaussian filter in Clampfit 10.2. A ValveLink 8.2 gravity flow Teflon perfusion system (AutoMate Scientific) was utilized to compare ionic current sizes generated by external monovalent or divalent ions.

## QUANTIFICATION AND STATISTICAL ANALYSES

Electrophysiology data gathered using pClamp10 (Molecular Devices) was imported into Origin-Pro 2017 (OriginLab Corporation) for statistical analyses. Kinetics of activation, inactivation, and deactivation were determined by fitting mono-exponential functions over the growing or decaying phases of each current trace. Electrophysiology data are shown as mean ± SEM where “n” refers to number of cells, provided in the figure legends, together with details of statistical tests used. Statistical significance between two groups was assessed by Student’s t-test, as stated. One-way ANOVA and the stated post hoc analysis was used for comparison of means between three or more groups.

## DATA AND CODE AVAILABILITY

PDB files of the AlphaFold2 generated molecular models of Cav3-12a and Cav3-12b channels for giant pond snail, *Lymnaea stagnalis* and nematode, *Caenorhabditis elegans* are provided as Supplementary Attachments 1 to 4.

### Supplementary information

PDB files of AlphaFold structures for molluscan (*Lymnaea stagnalis*) Cav3-12a and Cav3-12b and nematode (*Caenorhabditis elegans*) Cav3-12a and Cav3-12b

## Acknowledgments

This work was funded through a NSERC Discovery Grant and Heart and Stroke Foundation of Canada Grant-In-Aid to J.D.S, an NSERC grant to BSZ, and an NSERC Canada Graduate Scholarship to RFS. Computations were performed, in part, using AlphaFold2 software installed at Compute Canada (www.computecanada.ca).

## Author contributions

Conceived and designed the experiments, analyzed the data, and contributed reagents/materials/analysis tools: B.Z. and J.D.S. Performed the experiments: W.G., R.F.S., B.Z. and J.D.S. Wrote the paper: B.Z. and J.D.S.

## References

Bargmann, C. I. 1998. Neurobiology of the Caenorhabditis elegans genome. Science, 282, 2028–33.

Chakrabarti, N., Ing, C., Payandeh, J., Zheng, N., Catterall, W. A. & Pomes, R. 2013. Catalysis of Na+ permeation in the bacterial sodium channel Na(V)Ab. Proc Natl Acad Sci U S A, 110, 113–316.

Franks, C. J., Pemberton, D., Vinogradova, I., Cook, A., Walker, R. J. & Holden-Dye, L. 2002. Ionic basis of the resting membrane potential and action potential in the pharyngeal muscle of Caenorhabditis elegans. J Neurophysiol, 87, 954–61.

Fux, J. E., MEhta, A., Moffat, J. & Spafford, J. D. 2018. Eukaryotic Voltage-Gated Sodium Channels: On Their Origins, Asymmetries, Losses, Diversification and Adaptations. Front Physiol, 9, 1406.

Guan, W., Stephens, R. F., Mourad, O., Mehta, A., Fux, J. & Spafford, J. D. 2020. Unique cysteine-enriched, D2L5 and D4L6 extracellular loops in CaV3 T-type channels alter the passage and block of monovalent and divalent ions. Sci Rep, 10, 12404.

Heinemann, S. H., Terlau, H., Stuhmer, W., Imoto, K. & Numa, S. 1992. Calcium channel characteristics conferred on the sodium channel by single mutations. Nature, 356, 441–3.

Hille, B. 2001. Ion Channels of Excitable Membranes, Third Edition. In: Associates, S. (ed.). Sunderland, Mass.: Sinauer Associates.

Karmazinova, M., Beyl, S., Stary-Weinzinger, A., Suwattanasophon, C., Klugbauer, N., Hering, S. & Lacinova, L. 2010. Cysteines in the loop between IS5 and the pore helix of Ca(V)3.1 are essential for channel gating. Pflugers.Arch., 460, 1015–1028.

Schlief, T., Schonherr, R., Imoto, K. & Heinemann, S. H. 1996. Pore properties of rat brain II sodium channels mutated in the selectivity filter domain. Eur Biophys J, 25, 75–91.

Senatore, A., Boone, A. N. & Spafford, J. D. 2011. Optimized transfection strategy for expression and electrophysiological recording of recombinant voltage-gated ion channels in HEK-293T cells. J Vis Exp.

Senatore, A., Guan, W., Boone, A. N. & Spafford, J. D. 2014. T-type channels become highly permeable to sodium ions using an alternative extracellular turret region (S5-P) outside the selectivity filter. J Biol Chem, 289, 11952–69.

Senatore, A. & Spafford, J. D. 2010. Transient and big are key features of an invertebrate T-type channel (LCav3) from the central nervous system of Lymnaea stagnalis. J Biol Chem, 285, 744–758.

Steger, K. A., Shtonda, B. B., Thacker, C., Snutch, T. P. & Avery, L. 2005. The C. elegans T-type calcium channel CCA-1 boosts neuromuscular transmission. J Exp Biol, 208, 2191–2203.

Stephens, R. F., Guan, W., Zhorov, B. S. & Spafford, J. D. 2015. Selectivity filters and cysteine-rich extracellular loops in voltage-gated sodium, calcium, and NALCN channels. Front Physiol, 6, 153.

Tang, S., Mikala, G., Bahinski, A., Yatani, A., Varadi, G. & Schwartz, A. 1993. Molecular localization of ion selectivity sites within the pore of a human L-type cardiac calcium channel. J Biol Chem, 268, 13026–9.

Todorovic, S. M., Jevtovic-Todorovic, V., Meyenburg, A., Mennerick, S., Perez-Reyes, E., Romano, C., Olney, J. W. & Zorumski, C. F. 2001. Redox modulation of T-type calcium channels in rat peripheral nociceptors. Neuron, 31, 75–85.

Vinogradova, I., Cook, A. & Holden-Dye, L. 2006. The ionic dependence of voltage-activated inward currents in the pharyngeal muscle of Caenorhabditis elegans. Invert Neurosci, 6, 57–68.

Wang, D. W., George, A. L., JR. & Bennett, P. B. 1996. Comparison of heterologously expressed human cardiac and skeletal muscle sodium channels. Biophys J, 70, 238–45.

Zhao, Y., Huang, G., Wu, Q., Wu, K., Li, R., Lei, J., Pan, X. & Yan, N. 2019. Cryo-EM structures of apo and antagonist-bound human Cav3.1. Nature, 576, 492–497.

